## Supplementary Information for "Contextual AI models for single-cell protein biology"

### Contents

**S1 Supplementary Notes** **S3**
S1.1 Supplementary Note S1: Analyzing PINNACLE's embeddings for proteins with similar function . . . . . S3
S1.2 Supplementary Note S2: Benchmarking contextual 3D structure-based protein representations S4
S1.3 Supplementary Note S3: Extending PINNACLE to generate cell-level embeddings . . . . . S5

**S2 Supplementary Figures** **S6**

**S3 Supplementary Tables** **S17**

**S4 Supplementary References** **S21**

### S1 Supplementary Notes

#### S1.1 Supplementary Note S1: Analyzing PINNACLE's embeddings for proteins with similar function

**Dataset.** We extract human housekeeping genes from the Housekeeping and Reference Transcript Atlas (https://housekeeping.unicamp.br/)[1] and marker genes from the human gold standard T lymphocyte-specific protein functional networks from HumanBase (https://hb.flatironinstitute.org/) (accessed on November 20th, 2023)[2]. From HumanBase, only edges of level C1 (i.e., tissue-specific) are kept. The nodes corresponding to these edges are considered to be marker genes for cell types in the family of T lymphocytes. The lists of marker and housekeeping genes do not overlap, as we remove any overlapping housekeeping genes from the list of marker genes.

**Analysis.** We compare embedding similarities of a marker (orange) or housekeeping (gray) gene's contextualized protein representation (from PINNACLE) across different cell type contexts. For each marker or housekeeping gene, its cell type-specific protein representations are compared in similar contexts (i.e., between different T lymphocyte cell types; a total of 10 T lymphocyte cell types) or different contexts (i.e., between a T lymphocyte cell type and a non-immune cell type; a total of 115 non-immune cell types). We perform the two-sample Kolmogorov-Smirnov test (via `ks_2samp` from `scipy`).

**Results.** Although PINNACLE learns protein representations using context-aware protein, cell type, and tissue networks alone, it effectively captures protein functions. We analyze the embedding similarities of contextualized protein representations for marker and housekeeping genes across cell type contexts. For each T lymphocyte marker or housekeeping gene, we compare its cell type-specific protein representations in similar contexts (i.e., between different T lymphocyte cell types) and in different contexts (i.e., between a T lymphocyte cell type and a non-immune cell type). Housekeeping genes exhibit higher embedding similarity in similar contexts than marker genes (Supplementary Figure S5; $p$-value $= 3.2 \times 10^{-14}$). This result aligns with the expectation that housekeeping genes maintain shared functions across these cell types. Housekeeping genes in different contexts also show higher embedding similarity than marker genes (Supplementary Figure S5; $p$-value $= 1.0 \times 10^{-91}$), reflecting their consistent functions across non-immune cell types. Conversely, marker genes in similar contexts display higher embedding similarity than those in different contexts (Supplementary Figure S5; $p$-value $= 3.1 \times 10^{-26}$), consistent with their specificity to T lymphocyte cell types. Their protein representations are more similar within T lymphocyte contexts compared to when these marker genes are in the context of non-immune cell types. These analyses suggest that the protein embedding regions in PINNACLE are organized according to cellular contexts, potentially capturing subtle nuances not explicitly included in the training dataset or the model itself. This encompasses the possibility of cell type-dependent roles for proteins, a complexity that can enhance our understanding of protein functions across different biological contexts. Such insights warrant further investigation into proteins with context-specific and non-specific functions.

#### S1.2 Supplementary Note S2: Benchmarking contextual 3D structure-based protein representations

**Results.** We benchmark our contextualized protein representations (structure-free) and contextualized structure-based protein representations against two null distributions and four context-free approaches. We show that randomly sampling pairs of proteins from different cell type contexts, padded (no 3D structure; score gap $-0.0431$) or concatenated with the structure-based protein representations (score gap $-0.0356$), cannot produce the score gap observed in the contextualized protein representations (PINNACLE without 3D structure) nor contextualized structure-based protein representations (PINNACLE with 3D structure) (Supplementary Figure S7). Similarly, context-free protein representations cannot predict intercellular communication (i.e., protein interactions between different cell types). Such is demonstrated using context-free protein representations generated by a graph attention neural network[3] on the global reference protein interaction network (i.e., GAT), padded (no 3D structure; score gap $-0.1319$) and concatenated with the structure-based protein representations (score gap $-0.0486$), and context-free protein representations generated by BIONIC[4], a graph convolutional neural network designed for multi-modal network integration, padded (score gap $0.0046$) and concatenated with the structure-based protein representations (score gap $0.0043$). Our benchmarking results suggest that incorporating context can improve 3D structure prediction of protein interactions.

##### S1.3 Supplementary Note S3: Extending PINNACLE to generate cell-level embeddings

Unlike approaches that generate cell embeddings to advance cell-level downstream tasks, such as batch correction and cell type annotation[5–7], PINNACLE generates protein representations across cell types for precise protein-level prediction at cell type resolution. PINNACLE learns embeddings of cell types and tissues as a means to inject cellular and tissue organization (via the metagraph) into the unified protein embedding space. To enable cell-level characterization, PINNACLE can easily be extended to learn cell (rather than cell *type*) embeddings.

#### S2 Supplementary Figures

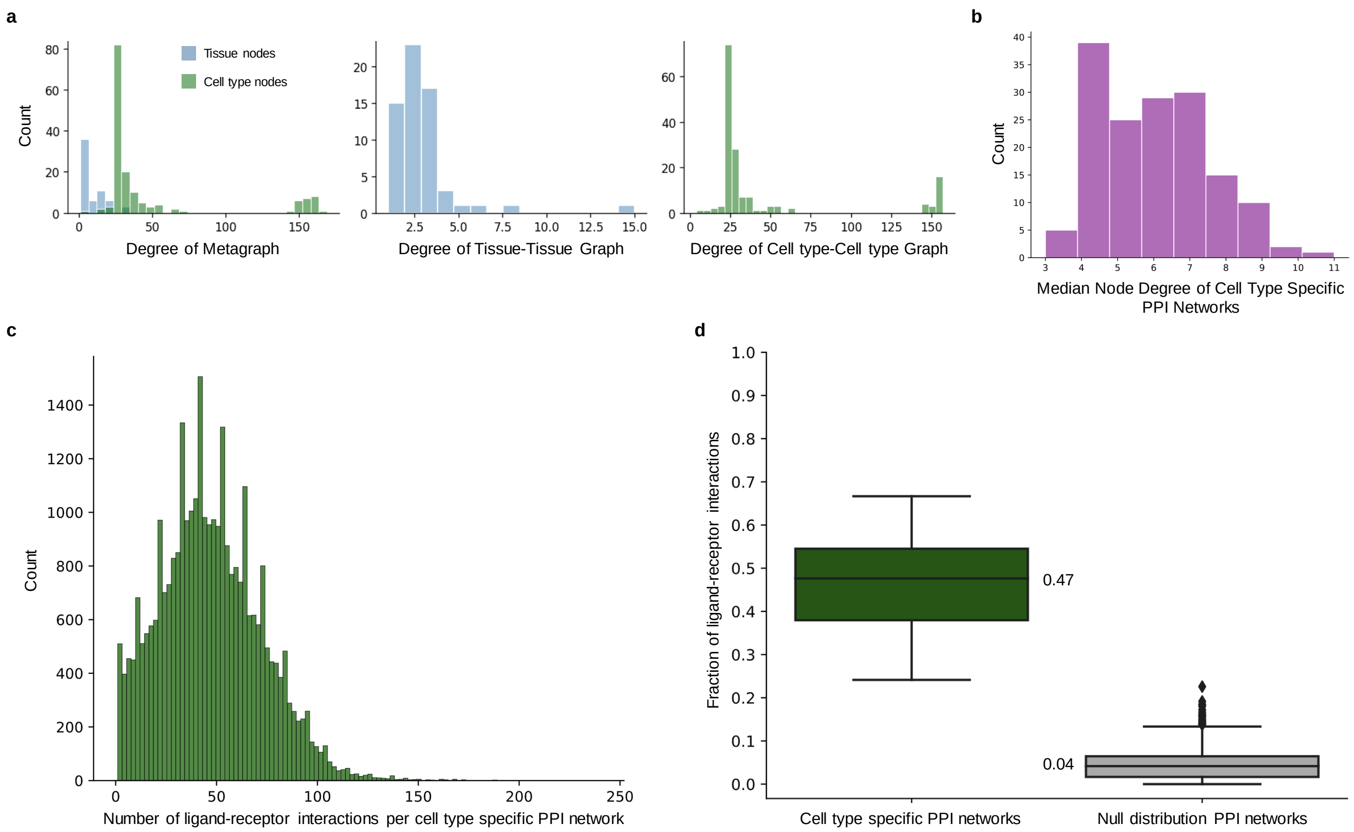


**Figure S1: Network properties of the metagraph and cell type specific protein interaction networks.** **(a-b)** Degree distributions of the metagraph and cell type specific protein interaction (PPI) networks. **(a)** Degree distributions of the metagraph (composed of cell type-cell type, cell type-tissue, and tissue-tissue edges), tissue-tissue graph, and cell type-cell type graph. The median, maximum, and minimum degrees for the metagraph are 24, 169, 1; for the tissue-tissue graph are 2, 15, 1; and for the cell type-cell type graph are 24, 157, 4. **(b)** Distribution of the median node degree of each cell type specific PPI network. The median, maximum, and minimum of median node degree across cell type specific PPI networks are 6, 11, and 3, respectively. **(c-d)** Enrichment analysis of ligand-receptor interactions in the cell type specific PPI networks. We utilize CellPhoneDB[8] to predict interactions between cell types in our metagraph by identifying significantly expressed ligand-receptor (LR) interactions between pairs of cell types in our dataset. **(c)** Shown is a histogram of the number of significant LR interactions per cell type specific PPI network predicted by CellPhoneDB. **(d)** We hypothesize that the predicted LR interactions are enriched in our cell type specific PPI networks. To quantify the enrichment of LR interactions, we calculate the fraction of LR interactions where the corresponding ligand and receptor proteins are activated in the cell type pair (i.e., for a LR interaction identified between cell types A and B, the ligand protein is activated in cell type A's PPI network and the receptor protein is activated in cell type B's PPI network). We compare the fraction of LR pairs that are activated in our cell type specific PPI networks against the fraction of LR pairs that are activated in null distribution PPI networks. For each cell type specific PPI network, we generate 100 null distribution PPI networks by sampling the same number of nodes with a similar degree distribution[9]. Degree distribution is preserved by binning nodes such that there are at least 100 nodes in each bin, and nodes are then randomly sampled within the appropriate degree interval[9]. We find that our cell type specific PPI networks have a significantly higher fraction of ligand-receptor pairs activated ($0.47 \pm 0.12$) than the null distribution PPI networks ($0.04 \pm 0.04$); $n =$ 2,020 pairs of cell type specific PPI networks, of which 20 are pairs of real cell type specific PPI networks and 2,000 are pairs of null cell type specific PPI networks. Note that the ligand-receptor interactions considered in both analyses are those where the genes corresponding to the ligands and receptors are known. However, this does not factor into our construction of the edges/interactions between cell types (CCI). The bounds of the box show the quartiles of the data, the center indicates the median value of the data, and the whiskers represent the farthest data point within $1.5 \times$ IQR.

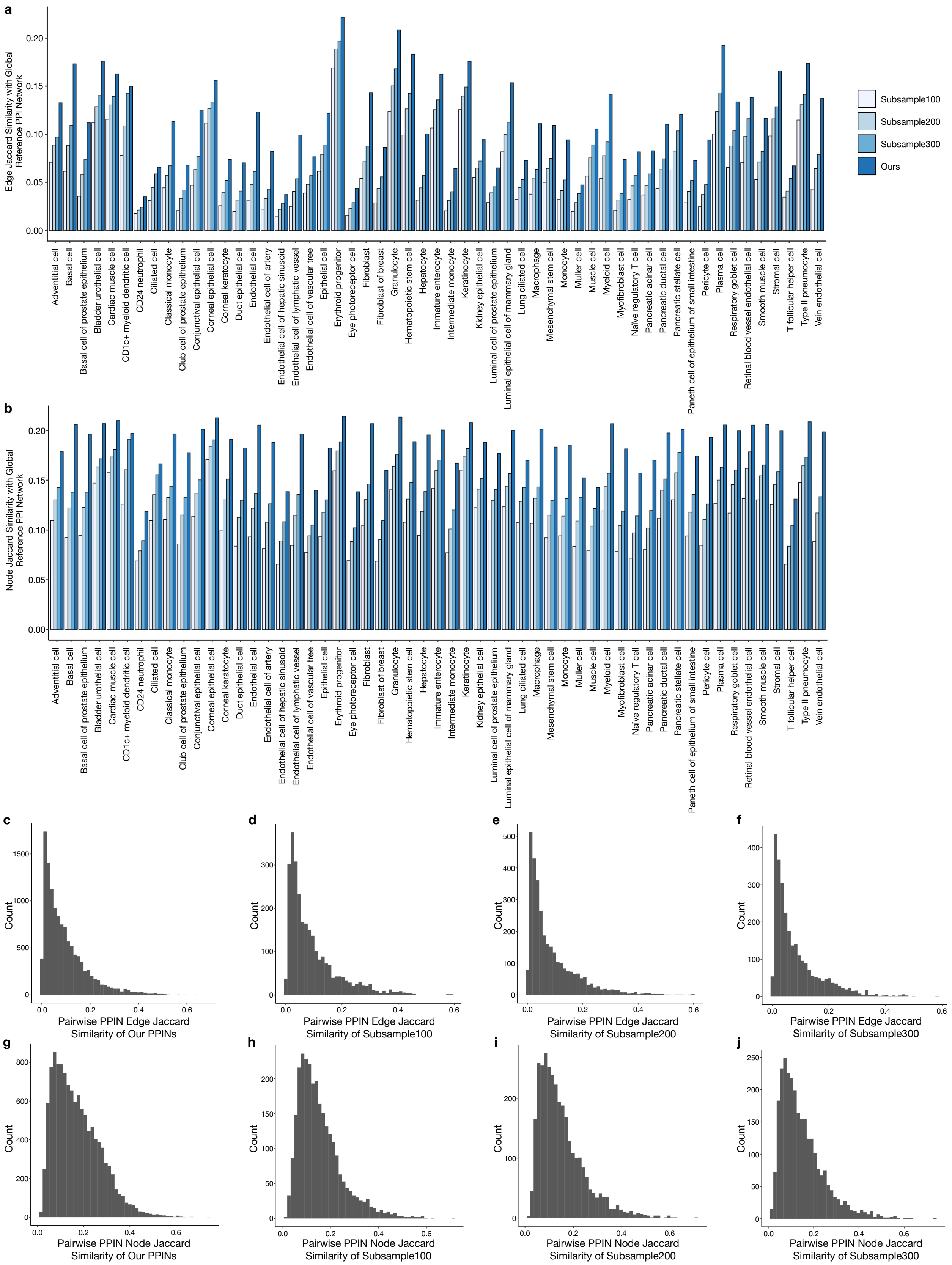


**Figure S2: Sensitivity analysis of network construction.** To examine whether cell types with fewer cells are poorly represented in our networks, we construct networks after subsampling equal numbers of cells per cell type. We compare our finalized networks (no subsampling of cells) against approaches that subsample 100, 200, and 300 cells. We find that our approach yields networks that are maximally similar to the global reference network yet maintain specificity to cell type context. **(a)** Edge and **(b)** node Jaccard similarity of a cell type specific PPIN to the global reference PPIN. **(c-j)** Distribution of edge jaccard similarity between PPINs constructed by **(c)** our finalized approach and subsampling **(d)** 100, **(e)** 200, and **(f)** 300 cells. **(g-j)** Distribution of node jaccard similarity between PPINs constructed by **(g)** our finalized approach and subsampling **(h)** 100, **(i)** 200, and **(j)** 300 cells.

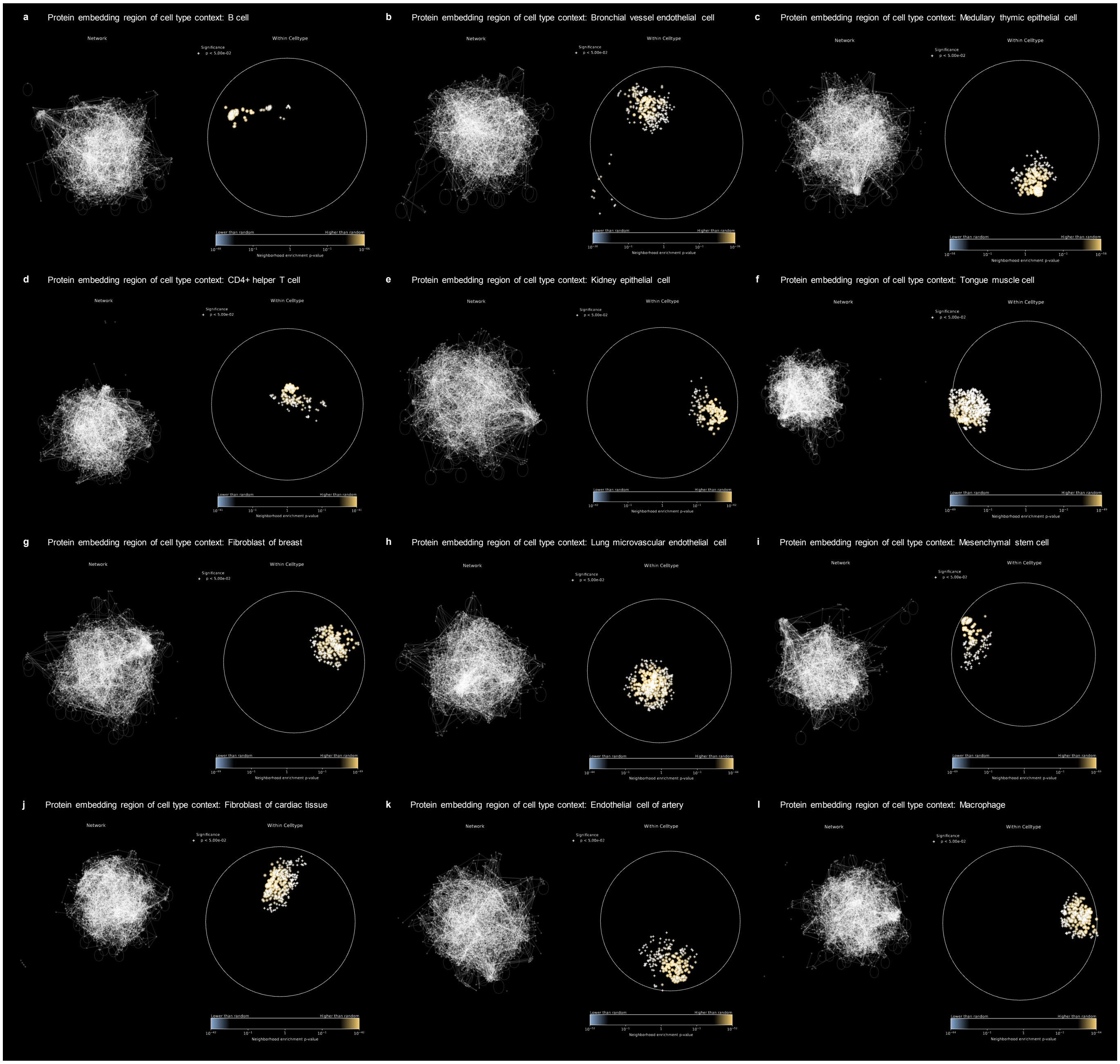


**Figure S3: Spatial enrichment analysis of PINNACLE's protein embedding regions.** For each cell type specific set of protein embeddings generated by PINNACLE, we sample a subset to construct a similarity network and perform spatial enrichment analysis using SAFE[10]. Shown for each cell type context is the network (left) and enrichment landscape (right). Dots represent the neighborhood enrichment $p$-value; crosses indicate a significant $p$-value $< 0.05$; hypergeometric test, adjusted using the Benjamin-Hochberg false discovery rate correction with significance cutoff $\alpha = 0.05$.

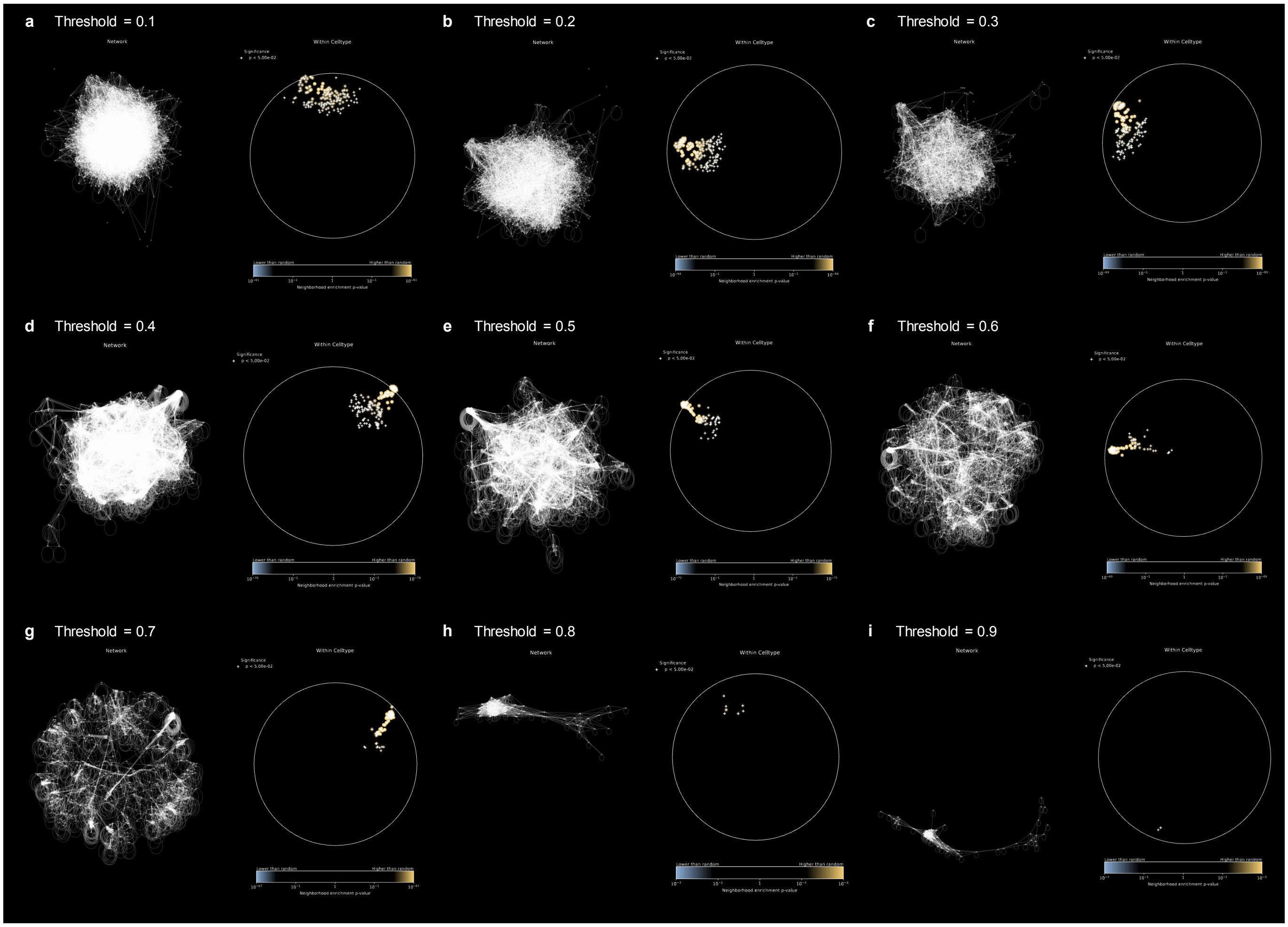


**Figure S4: Spatial enrichment analysis of PINNACLE's protein embedding regions across thresholds.** From the mesenchymal stem cell type specific protein embeddings generated by PINNACLE, we sample a subset to construct a similarity network and perform spatial enrichment analysis using SAFE[10]. Networks are constructed using a similarity threshold $t \in [0.1, 0.2, 0.3, 0.4, 0.5, 0.6, 0.7, 0.8, 0.9]$. Shown for each threshold is the network (left) and enrichment landscape (right). Dots represent the neighborhood enrichment $p$-value; crosses indicate a significant $p$-value $< 0.05$; hypergeometric test, adjusted using the Benjamin-Hochberg false discovery rate correction with significance cutoff $\alpha = 0.05$.

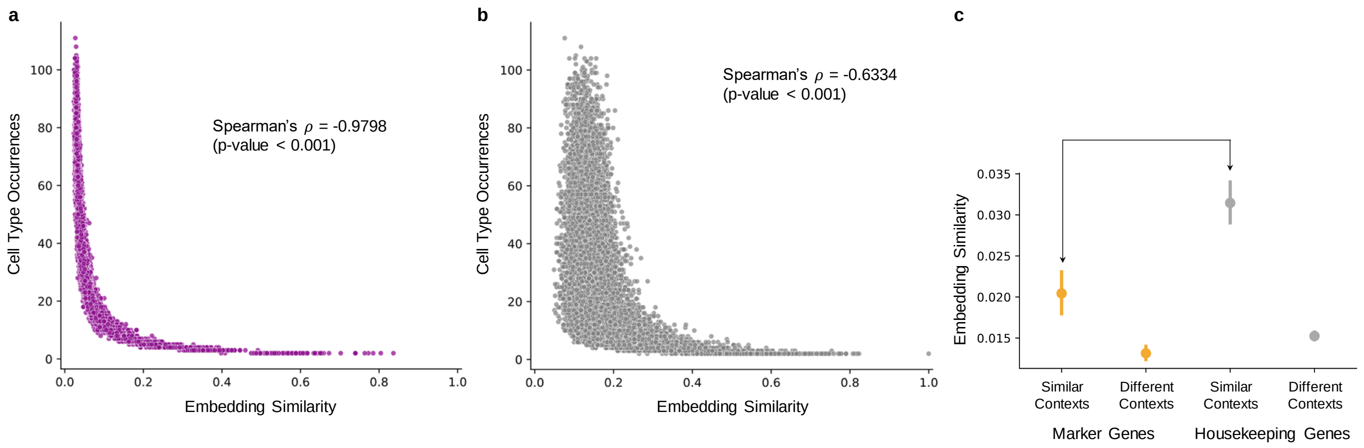


**Figure S5: Embedding similarity based on proteins' cell type activation and function.** **(a-b)** Each dot represents a protein that is activated in at least two cell types. Shown is the average cosine similarity of embeddings for each protein as a function of the number of cell types that it is activated in **(a)** with ($p$-value $< 0.001$) and **(b)** without ($p$-value $< 0.001$) cellular and tissue context. Both Spearman correlation statistical tests for **(a)** and **(b)** are two-sided. **(c)** Comparison of embedding similarities of a marker (orange) or housekeeping (gray) gene's contextualized protein representation (from PINNACLE) across different cell type contexts. The marker genes are specific to cell types in the family of T lymphocytes (a total of 10 T lymphocyte cell types). For each marker/housekeeping gene, its cell type specific protein representations are compared in similar contexts (i.e., between different T lymphocyte cell types) or different contexts (i.e., between a T lymphocyte cell type and a non-immune cell type; a total of 115 non-immune cell types). All comparisons between these four groups shown are statistically significant. Cosine embedding similarity is used to compare contextualized protein representations. Data are represented as mean values with error bars indicating a 95% confidence interval.

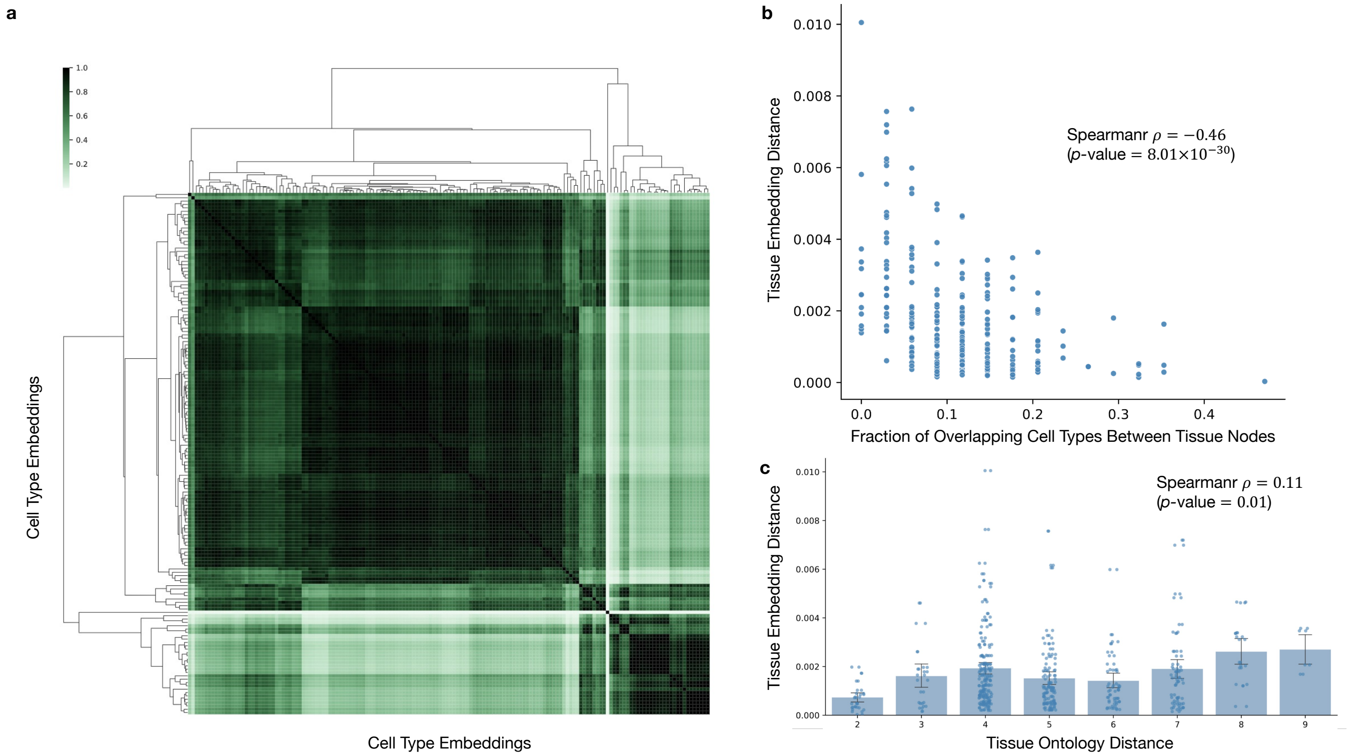


**Figure S6: Evaluation of PINNACLE's cell type and tissue representations.** **(a)** We quantify the quality of PINNACLE's cell type representations by calculating pairwise similarities of cell type representations. Pairwise similarities are computed via cosine similarity. We expect several major groups of cell type representations that are organized according to cellular and tissue hierarchy and acting as anchors for our complete set of cell type representations. This implies that the contextual information being transferred between the representations of cell types and proteins reflects the tissue hierarchy. Our results show that the local organization of PINNACLE's cell type representations (i.e., identity of cell types in each group) reflects cellular communication, and the global organization of cell type representations (i.e., proximity of groups to each other) reflects tissue organization. Since PINNACLE's protein representations are embedded near their corresponding cell type representation, such organization is enforced among the contextualized protein representations as well. **(b)** Correlation between cosine distance of tissue representations and the fraction of overlapping cell types neighbors between the tissue pair. Spearman $\rho = -0.46$ with $p$-value $= 8.01 \times 10^{-30}$. **(c)** Correlation between PINNACLE's tissue embedding distance to tissue ontology distance for leaf nodes in the metagraph. Spearman $\rho = 0.11$ with $p$-value $= 0.01$. All Spearman correlation statistical tests are two-sided. Data are represented as mean values with error bars indicating a 95% confidence interval. Both panels show $n = 548$ pairwise comparison calculations.

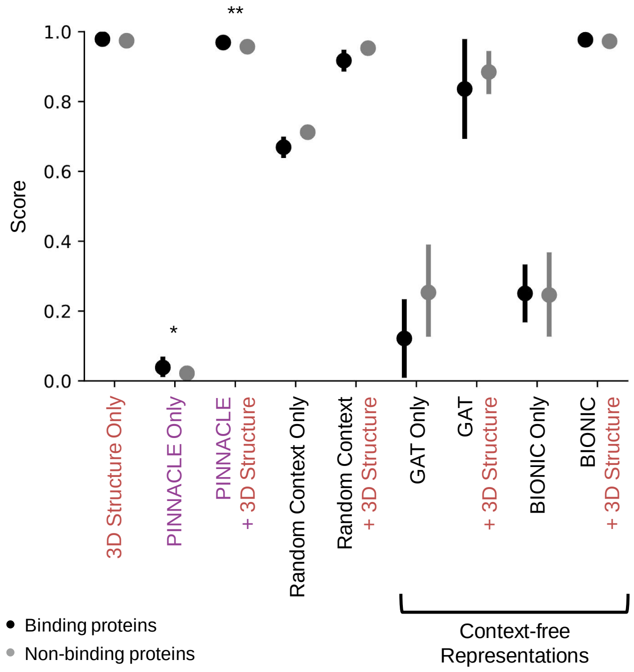


| Representation Type | Binding Proteins | Non-binding Proteins | Gap (p-value) |
|---|---|---|---|
| 3D Structure Only | 0.9789 ± 0.0004 | 0.9742 ± 0.0078 | 0.0047 (0.2121) |
| **PINNACLE Only** | **0.0385 ± 0.1531** | **0.0218 ± 0.1081** | **0.0167 (0.0299)** |
| **PINNACLE + 3D Structure** | **0.9690 ± 0.0049** | **0.9571 ± 0.0127** | **0.0119 ($< 10^{-5}$)** |
| Random Context Only | 0.6691 ± 0.1642 | 0.7122 ± 0.0445 | −0.0431 (1.0) |
| Random Context + 3D Structure | 0.9172 ± 0.1732 | 0.9529 ± 0.0277 | −0.0356 (1.0) |
| GAT Only | 0.1214 ± 0.1056 | 0.2533 ± 0.2898 | −0.1319 (0.6939) |
| GAT + 3D Structure | 0.8360 ± 0.1360 | 0.8847 ± 0.1255 | −0.0486 (0.5706) |
| BIONIC Only | 0.2506 ± 0.0760 | 0.2460 ± 0.2536 | 0.0046 (0.4556) |
| BIONIC + 3D Structure | 0.9769 ± 0.0018 | 0.9725 ± 0.0087 | 0.0043 (0.2797) |

**Figure S7: Benchmarking context-free and contextualized 3D structure protein representations.** Shown are binding and non-binding scores (i.e., cosine similarity) of proteins when using only 3D structure-based protein representations ($p$-value $= 0.2121$; $n = 22$ pairwise comparisons between 2 binding and 20 non-binding pairs), PINNACLE's contextualized protein representations (without 3D structural information; $p$-value $= 0.0299$; $n =$ 7,956 pairwise computations between 180 binding and 7,776 non-binding pairs), contextualized structure-based protein representations ($p$-value $< 10^{-5}$; $n =$ 7,956 pairwise computations between 180 binding and 7,776 non-binding pairs), and baseline models. The baseline models are random context only (i.e., randomly sampling pairs of PINNACLE's protein representations from different cell type contexts; $p$-value $= 1.0$; $n =$ 7,956 pairwise computations between 180 "binding" and 7,776 "non-binding" pairs), concatenating random context protein representations with 3D structure-based protein representations ($p$-value $= 1.0$; $n =$ 7,956 pairwise computations between 180 "binding" and 7,776 "non-binding" pairs), GAT only (i.e., context-free protein representations generated by a graph attention neural network[3] on the global reference interactome; $p$-value $= 0.6939$; $n = 22$ pairwise comparisons between 2 binding and 20 non-binding pairs), concatenating GAT protein representations with 3D structure-based protein representations ($p$-value $= 0.5706$; $n = 22$ pairwise comparisons between 2 binding and 20 non-binding pairs), BIONIC only (i.e., context-free protein representations generated by BIONIC[4], a graph convolutional neural network designed for multi-modal network integration; $p$-value $= 0.4556$; $n = 22$ pairwise comparisons between 2 binding and 20 non-binding pairs), and concatenating BIONIC protein representations with 3D structure-based protein representations ($p$-value $= 0.2797$; $n = 22$ pairwise comparisons between 2 binding and 20 non-binding pairs). Note that all protein representations have consistent dimensions (328 = 200 structure-based protein representation + 128 context-aware/-free protein representation) to ensure that they are comparable. The protein representations without 3D structure are padded with 0's (i.e., null 3D structure-based protein representation). The significance of the score gaps between binding and non-binding proteins is measured using a one-sided non-parametric permutation test. Data are represented as mean values with error bars indicating a 95% confidence interval.

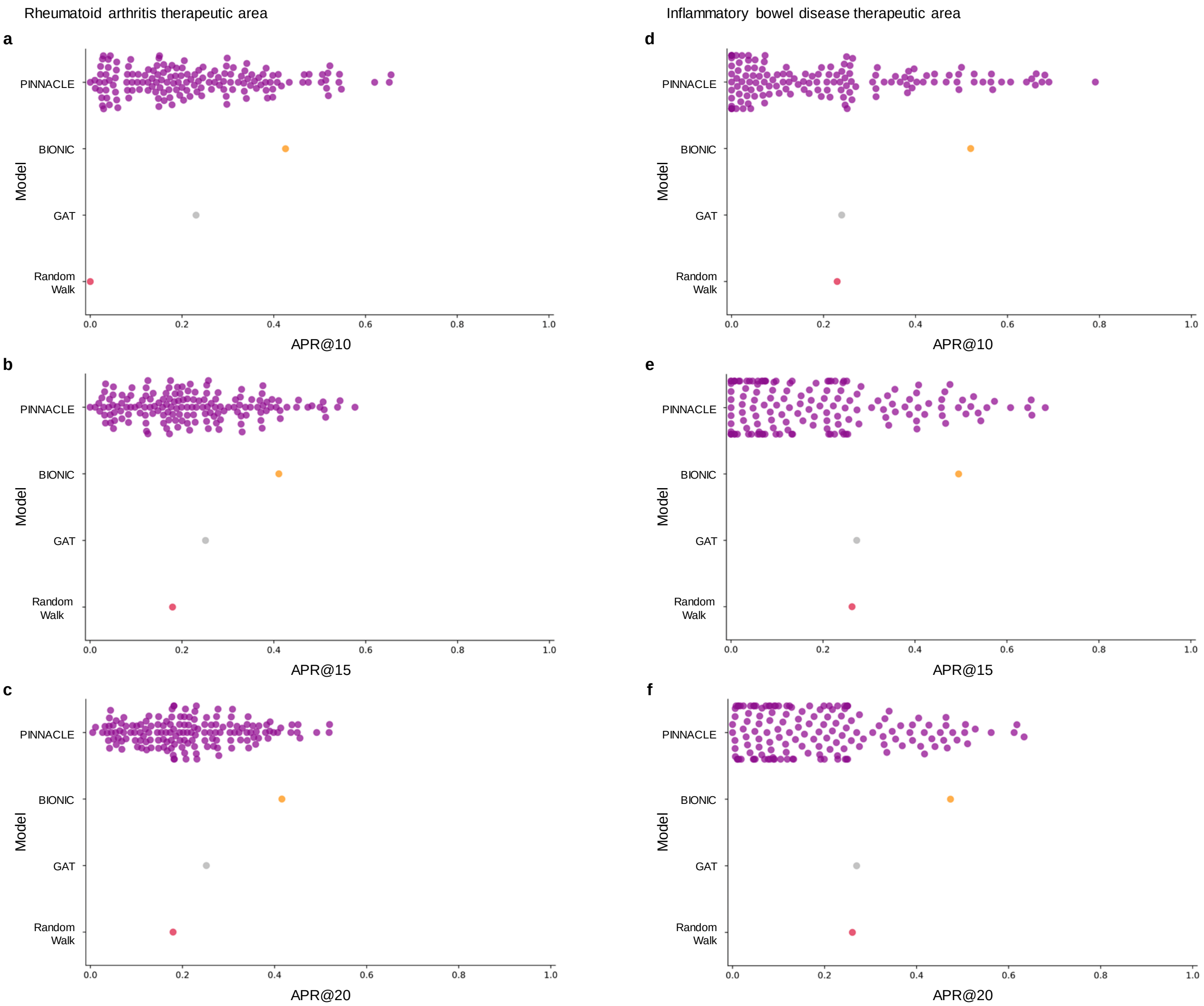


**Figure S8: Performance of therapeutic target prioritization models for rheumatoid arthritis and inflammatory bowel diseases.** Benchmarking of context-aware and context-free approaches for **(a-c)** RA and **(d-f)** IBD therapeutic areas. Each dot is the performance (averaged across 10 random seeds) of protein representations from a given context (i.e., cell type context for PINNACLE, context-free global reference protein interaction network for GAT and random walk, and context-free multi-modal protein interaction network for BIONIC). In the model for the RA therapeutic area: **(a)** at APR@10, 100% of cell types (156 out of 156) outperform the random walk model, 44.2% of cell types (69 out of 156) outperform GAT, and 11.5% of cell types (18 out of 156) outperform BIONIC. **(b)** At APR@15, 58.3% (91 out of 156) outperform the random walk model, 38.5% of cell types (60 out of 156) outperform GAT, and 9.0% of cell types (14 out of 156) outperform BIONIC. **(c)** At APR@20, 59.0 (92 out of 156) outperform the random walk model, 34.6% of cell types (54 out of 156) outperform GAT, and 5.1% of cell types (8 out of 156) outperform BIONIC. In the model for the IBD therapeutic area: **(d)** at APR@10, 39.5% (60 out of 152) outperform the random walk model, 38.2% of cell types (58 out of 152) outperform GAT, and 10.5% of cell type (16 out of 152) outperform BIONIC. **(e)** At APR@15, 28.3% (43 out of 152) outperform the random walk model and GAT, and 8.6% of cell types (13 out of 152) outperform BIONIC. **(f)** At APR@20, 26.3% (40 out of 152) outperform the random walk model and GAT, and 6.6% of cell types (10 out of 152) outperform BIONIC.

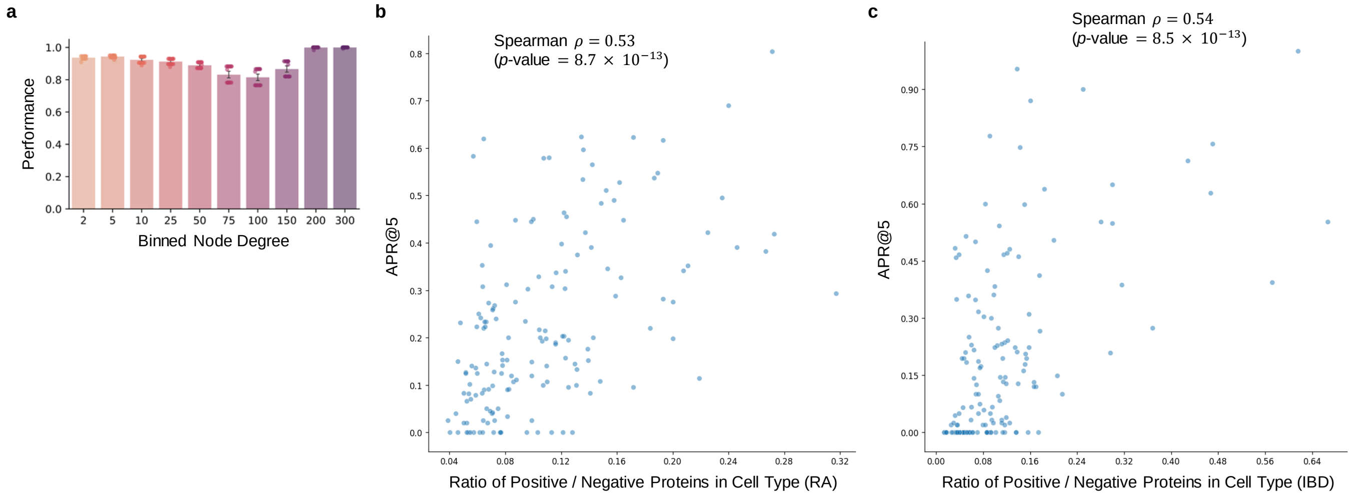


**Figure S9: Correlating downstream performance on rheumatoid arthritis and inflammatory bowel diseases with protein degree and network enrichment. (a)** Correlation between the node degrees of proteins (in the cell type specific protein interaction networks) and the downstream performance of their learned representations. Combining the RA and IBD prediction results, the Spearman $\rho = 0.087$ with $p$-value $= 0.223$ ($n =$ 36,229, consisting of 3,165 positive protein examples with label $y = 1$ and 33,064 negative protein examples with label $y = 0$). For RA only, the Spearman $\rho = 0.205$ with $p$-value $= 0.041$ ($n =$ 26,773, consisting of 2,382 positive protein examples with label $y = 1$ and 24,391 negative protein examples with label $y = 0$). For IBD only, the Spearman $\rho = 0.024$ with $p$-value $= 0.810$ ($n =$ 9,456, consisting of 783 positive protein examples with label $y = 1$ and 8,673 negative protein examples with label $y = 0$). Data are represented as mean values with error bars indicating a 95% confidence interval. **(b-c)** Correlation between PINNACLE's performance and network enrichment. **(b)** Comparing PINNACLE's predicted performance (APR@5) and the ratio of positive to negative proteins in each cell type for RA (Spearman $\rho = 0.53$ with $p$-value $= 8.7 \times 10^{-13}$; $n =$ 26,773, consisting of 2,382 positive proteins with label $y = 1$ and 24,391 negative proteins with label $y = 0$). **(c)** Comparing PINNACLE's predicted performance (APR@5) and the ratio of positive to negative proteins in each cell type for IBD (Spearman $\rho = 0.54$ with $p$-value $= 8.5 \times 10^{-13}$; $n =$ 9,456, consisting of 783 positive proteins with label $y = 1$ and 8,673 negative proteins with label $y = 0$). All Spearman correlation statistical tests are two-sided.

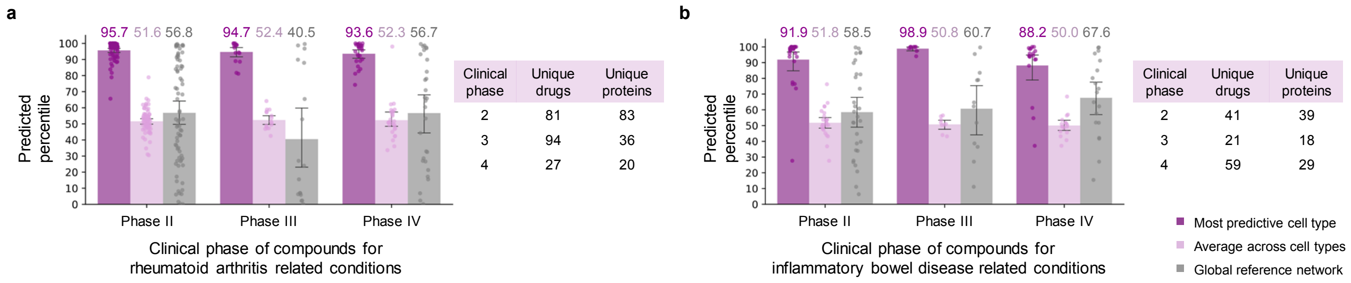


**Figure S10: Performance of therapeutic target prioritization models for rheumatoid arthritis and inflammatory bowel diseases stratified by clinical trials.** Comparison of the percentiles of drug targets across cell types, in their best-performing cell types, and in the context-free global reference model, stratified by clinical phase of compounds for **(a)** RA and **(b)** IBD. The table shows the number of unique drugs in each clinical phase, as well as the numbers of unique proteins targeted by those drugs. Data are represented as mean values with error bars indicating a 95% confidence interval.

### S3 Supplementary Tables

**Table S1: Predicted ranks of relevant cell types for rheumatoid arthritis.** Shown are predicted ranks of subtypes of T cells, natural killer (NK) cells, dendritic cells, B cells, monocytes, and myeloid cells. These cell types have been demonstrated in existing literature to be involved with rheumatoid arthritis[11–13].

| **Cell type** | **Subtypes** | **PINNACLE-Predicted Rank** |
|---|---|---|
| T cell | CD4+ helper T cell | 1 |
| | CD4+ $\alpha\beta$ memory T cell | 2 |
| | Regulatory T cell | 6 |
| | CD8+ $\alpha\beta$ cytotoxic T cell | 27 |
| | DN1 thymic pro-T cell | 33 |
| | Mature natural killer T cell | 34 |
| | Naive regulatory T cell | 39 |
| | Type I natural killer T cell | 49 |
| | Naive thymus-derived CD4+ $\alpha\beta$ T cell | 77 |
| | CD8+ $\alpha\beta$ cytokine secreting effector T cell | 104 |
| Dendritic cell | CD1c+ myeloid dendritic cell | 3 |
| | CD141+ myeloid dendritic cell | 11 |
| | Dendritic cell | 37 |
| | Mature conventional dendritic cell | 38 |
| | Myeloid dendritic cell | 42 |
| | Plasmacytoid dendritic cell | 43 |
| | Liver dendritic cell | 50 |
| B cell | Memory B cell | 12 |
| | B cell | 29 |
| Natural killer cell | Natural killer cell | 17 |
| | Immature natural killer cell | 113 |
| Monocyte | Intermediate monocytes | 18 |
| | Non-classical monocytes | 68 |
| | Monocyte | 85 |
| | Classical monocytes | 91 |
| Myeloid cell | Myeloid progenitor | 71 |
| | Myeloid cell | 95 |

**Table S2: Predicted ranks of relevant cell types for inflammatory bowel diseases.** Shown are predicted ranks of subtypes of T cell, fibroblast, goblet cell, enterocyte, monocyte, natural killer cell, B cell, glial cell, dendritic cell, and macrophage. These cell types have been demonstrated in existing literature to be involved with inflammatory bowel diseases[14,15].

| Cell type | Subtypes | PINNACLE-Predicted Rank |
|---|---|---|
| T cell | CD4+ $\alpha\beta$ memory T cell | 1 |
| | Naive thymus-derived CD4+ $\alpha\beta$ T cell | 11 |
| | Regulatory T cell | 14 |
| | DN1 thymic pro-T cell | 15 |
| | Mature natural killer T cell | 16 |
| | CD8+ $\alpha\beta$ cytokine secreting effector T cell | 19 |
| | Type I natural killer T cell | 30 |
| | CD4+ helper T cell | 41 |
| | Naive regulatory T cell | 43 |
| | CD8+ $\alpha\beta$ cytotoxic T cell | 56 |
| Enterocyte | Enterocyte of epithelium of large intestine | 2 |
| | Mature enterocyte | 22 |
| | Immature enterocyte | 90 |
| | Enterocyte of epithelium of small intestine | 94 |
| | Intestinal enteroendocrine cell | 138 |
| Dendritic cell | Myeloid dendritic cell | 5 |
| | Dendritic cell | 10 |
| | CD1c+ myeloid dendritic cell | 13 |
| | CD141+ myeloid dendritic cell | 50 |
| | Liver dendritic cell | 54 |
| | Mature conventional dendritic cell | 67 |
| | Plasmacytoid dendritic cell | 72 |
| Goblet cell | Large intestine goblet cell | 7 |
| | Goblet cell | 33 |
| | Small intestine goblet cell | 75 |
| | Respiratory goblet cell | 109 |
| | Tracheal goblet cell | 110 |
| B cell | B cell | 9 |
| | Memory B cell | 45 |
| Monocyte | Intermediate monocyte | 26 |
| | Non-classical monocyte | 81 |
| | Monocyte | 89 |
| | Classical monocyte | 93 |
| Glial cell | Microglial cell | 29 |
| | Radial glial cell | 107 |
| Natural killer cell | Immature natural killer cell | 31 |
| | Natural killer cell | 73 |
| Fibroblast | Fibroblast | 49 |
| Macrophage | Macrophage | 74 |

**Table S3: Metagraph network statistics with different cutoff selection.** Data statistics of the metagraph with different cutoffs for the minimum number of significant ligand-receptor interactions between a pair of cell types to create an edge. Shown are the number of nodes, number of edges, and the average degree of the cell type-cell type interaction (CCI) graph and the metagraph (includes cell type-cell type, cell type-tissue, and tissue-tissue edges).

| Cutoff for Significant LRs | Component of Metagraph | Number of Nodes | Number of Edges | Average Degree |
|---|---|---|---|---|
| Cutoff = 1 (Original) | CCI Graph | 156 | 3,567 | 45.7 |
| | Metagraph (All) | 218 | 4,018 | 36.9 |
| Cutoff = 2 | CCI Graph | 156 | 1,808 | 23.2 |
| | Metagraph (All) | 218 | 2,259 | 20.7 |
| Cutoff = 3 | CCI Graph | 156 | 1,736 | 22.3 |
| | Metagraph (All) | 218 | 2,187 | 20.1 |
| Cutoff = 4 | CCI Graph | 156 | 1,640 | 21.0 |
| | Metagraph (All) | 218 | 2,091 | 19.2 |
| Cutoff = 5 | CCI Graph | 156 | 1,576 | 20.2 |
| | Metagraph (All) | 218 | 2,027 | 18.6 |

**Table S4: Sensitivity analysis of cutoff selection.** Sensitivity analysis to examine the impact of the cutoff value for the minimum required number of significant ligand-receptor interactions in the cell-type-to-cell-type graph on PINNACLE's embedding space. The first row consists of results from the complete model. The remaining four rows show results from cutoff values 2, 3, 4, and 5. All Spearman correlation statistical tests are two-sided.

| Model | Tissue Embedding Distance vs. Ontology Distance | Tissue Embedding Distance vs. Fraction of Cell Type Overlap (leaves only) |
|---|---|---|
| Complete model (Cutoff = 1) | Spearman $\rho = 0.36$<br>$p$-value $= 4.6 \times 10^{-119}$ | Spearman $\rho = -0.46$<br>$p$-value $= 8.01 \times 10^{-30}$ |
| Cutoff = 2 | Spearman $\rho = 0.21$<br>$p$-value $= 1.0 \times 10^{-37}$ | Spearman $\rho = -0.31$<br>$p$-value $= 2.4 \times 10^{-13}$ |
| Cutoff = 3 | Spearman $\rho = 0.22$<br>$p$-value $= 1.7 \times 10^{-41}$ | Spearman $\rho = -0.31$<br>$p$-value $= 7.8 \times 10^{-14}$ |
| Cutoff = 4 | Spearman $\rho = 0.25$<br>$p$-value $= 1.4 \times 10^{-53}$ | Spearman $\rho = -0.29$<br>$p$-value $= 2.3 \times 10^{-12}$ |
| Cutoff = 5 | Spearman $\rho = 0.38$<br>$p$-value $= 8.8 \times 10^{-129}$ | Spearman $\rho = -0.25$<br>$p$-value $= 3.7 \times 10^{-9}$ |

**Table S5: Ablation studies to interrogate the contribution of the metagraph.** The first row consists of results from the complete model. The remaining three rows show results from three types of ablations: removing cell-type-to-cell-type relationships (i.e., shuffling the cell type nodes' identities), removing tissue-to-tissue relationships (i.e., shuffling the tissue nodes' identities), and removing the metagraph (i.e., setting the weight of the metagraph-related terms in the loss function to zero). The performance metrics evaluate the models' ability to capture cell type and tissue organization in the embedding space. The second column is the correlation between tissue embedding distance (computed using the model's tissue representations) and tissue ontology distance; we expect a positive correlation. The third column is the correlation between tissue embedding distance and tissue ontology distance among the tissue leaf nodes of the metagraph; we expect a positive correlation. The fourth column is the correlation between tissue embedding distance and fraction of overlapping cell types; we expect a strong negative correlation. All Spearman correlation statistical tests are two-sided.

| **Model** | **Tissue Embedding Distance vs. Ontology Distance** | **Tissue Embedding Distance vs. Ontology Distance (leaves only)** | **Tissue Embedding Distance vs. Fraction of Cell Type Overlap (leaves only)** |
|---|---|---|---|
| Complete model | Spearman $\rho = 0.36$<br>$p$-value $= 4.6 \times 10^{-119}$ | Spearman $\rho = 0.11$<br>$p$-value $= 0.01$ | Spearman $\rho = -0.46$<br>$p$-value $= 8.01 \times 10^{-30}$ |
| Drop cell type-cell type graph | Spearman $\rho = 0.38$<br>$p$-value $= 2.3 \times 10^{-132}$ | Spearman $\rho = 0.10$<br>$p$-value $= 0.02$ | Spearman $\rho = -0.21$<br>$p$-value $= 4.25 \times 10^{-7}$ |
| Drop tissue-tissue graph | Spearman $\rho = -0.13$<br>$p$-value $= 1.2 \times 10^{-14}$ | Spearman $\rho = -0.15$<br>$p$-value $= 6.5 \times 10^{-4}$ | Spearman $\rho = -0.16$<br>$p$-value $= 2.5 \times 10^{-4}$ |
| Drop metagraph loss | Spearman $\rho = 0.30$<br>$p$-value $= 4.6 \times 10^{-79}$ | Spearman $\rho = -0.10$<br>$p$-value $= 0.02$ | Spearman $\rho = -0.19$<br>$p$-value $= 1.1 \times 10^{-5}$ |

**Table S6: Data split of downstream tasks.** Sizes of the train, validation, and test datasets for the rheumatoid arthritis (RA) PINNACLE model and inflammatory bowel disease (IBD) PINNACLE model. The numeric value outside the parentheses represents the number of protein representations across cell type contexts, and the numeric value inside the parentheses represents the number of unique protein identities. The numbers represent both positive (label $= 1$) and negative (label $= 0$) proteins. Note that the validation dataset set is sampled from the train dataset, which is fixed, at each run of the model. The numbers for train and validation datasets (columns 3-4) are from seed 1.

| **Dataset** | **Type of protein target** | **Proteins in train dataset (unique)** | **Proteins in validation dataset (unique)** | **Proteins in test dataset (unique)** |
|---|---|---|---|---|
| RA | Total | 17,408 (600) | 6,647 (195) | 25,137 (818) |
| | Positive | 1,319 (53) | 570 (21) | 2,226 (78) |
| | Negative | 16,089 (547) | 6,077 (174) | 22,911 (740) |
| IBD | Total | 27,652 (896) | 9,363 (297) | 8,864 (294) |
| | Positive | 1,210 (62) | 673 (26) | 731 (26) |
| | Negative | 26,442 (834) | 8,690 (271) | 8,133 (268) |

#### S4 Supplementary References


1. Hounkpe, B. W., Chenou, F., de Lima, F. & De Paula, E. V. HRT Atlas v1. 0 database: redefining human and mouse housekeeping genes and candidate reference transcripts by mining massive RNA-seq datasets. *Nucleic Acids Research* **49**, D947–D955 (2021).

2. Greene, C. S. *et al.* Understanding multicellular function and disease with human tissue-specific networks. *Nature Genetics* **47**, 569–576 (2015).

3. Brody, S., Alon, U. & Yahav, E. How attentive are graph attention networks? *ICLR* (2022).

4. Forster, D. T. *et al.* BIONIC: biological network integration using convolutions. *Nature Methods* **19**, 1250–1261 (2022).

5. Chen, H., Ryu, J., Vinyard, M. E., Lerer, A. & Pinello, L. SIMBA: single-cell embedding along with features. *Nature Methods* 1–11 (2023).

6. Theodoris, C. V. *et al.* Transfer learning enables predictions in network biology. *Nature* **618**, 616–624 (2023).

7. Cui, H. *et al.* scGPT: toward building a foundation model for single-cell multi-omics using generative AI. *Nature Methods* 1–11 (2024).

8. Efremova, M., Vento-Tormo, M., Teichmann, S. A. & Vento-Tormo, R. CellPhoneDB: inferring cell–cell communication from combined expression of multi-subunit ligand–receptor complexes. *Nature Protocols* **15**, 1484–1506 (2020).

9. Guney, E., Menche, J., Vidal, M. & Barábasi, A.-L. Network-based in silico drug efficacy screening. *Nature Communications* **7**, 10331 (2016).

10. Baryshnikova, A. Systematic functional annotation and visualization of biological networks. *Cell Systems* **2**, 412–421 (2016).

11. Lewis, M. J. *et al.* Molecular portraits of early rheumatoid arthritis identify clinical and treatment response phenotypes. *Cell Reports* **28**, 2455–2470 (2019).

12. Zhang, F. *et al.* Deconstruction of rheumatoid arthritis synovium defines inflammatory subtypes. *Nature* **623**, 616–624 (2023).

13. Vickovic, S. *et al.* Three-dimensional spatial transcriptomics uncovers cell type localizations in the human rheumatoid arthritis synovium. *Communications Biology* **5**, 129 (2022).

14. Smillie, C. S. *et al.* Intra-and inter-cellular rewiring of the human colon during ulcerative colitis. *Cell* **178**, 714–730 (2019).

15. Kong, L. *et al.* The landscape of immune dysregulation in Crohn's disease revealed through single-cell transcriptomic profiling in the ileum and colon. *Immunity* **56**, 444–458 (2023).
